# Tiled PCR amplification-based Whole Genome Sequencing and Phylogenetic Classification Accelerate the Implementation of Respiratory Syncytial Virus Genomic surveillance in Canada as a Pilot Study

**DOI:** 10.1101/2024.11.22.624816

**Authors:** Ruimin Gao, Cody Buchanan, Kerry Dust, Paul Van Caeseele, Henry Wong, Calvin Sjaarda, Prameet M. Sheth, Agatha N. Jassem, Jessica Minion, Nathalie Bastien

**Author notes:** **Co-Correspondence:** Ruimin Gao; Nathalie Bastien.

## Abstract

Whole genome sequencing (WGS) has emerged as a powerful tool to facilitate the study of existing and emerging infectious diseases. WGS-based genomic surveillance provides information on the genetic diversity and tracks the evolution of important viral pathogens including Respiratory Syncytial Virus (RSV). Development and implementation of robust tiled multiplex PCR amplification-based WGS assays will facilitate high-throughput RSV surveillance initiatives. In this study, we developed multiplex PCR assays for targeted enrichment of viral genomes using PrimalScheme (http://primal.zibraproject.org) to amplify over 97% of the genome in the majority of contemporaneous specimens tested. A pilot dataset comprising 52 RSVA and 37 RSVB genomes derived from Canadian clinical specimens during the 2022-2023 respiratory virus season were used to perform phylogenetic analyses using both near complete genome and Glycoprotein (G) sequences. Overall, the RSV phylogenetic tree built with whole genomes showed identical lineage clusters as that compared to the G gene, but showed more confidence and discriminatory features within individual lineage. Moreover, availability of whole genomes enabled the identification of a broader range of mutations, for instance the identified S377N, K272M, S276N, S211N, S206I and S209Q in Canadian fusion proteins that could be potentially associated with effectiveness of vaccines or antiviral-based therapeutics. In conclusion, the tiled-PCR amplification assays described offer a more streamlined approach to facilitate high-throughput, high sensitivity of RSV WGS, which is capable of supporting enhanced genomic surveillance initiatives, as well as the more comprehensive genomic analyses required to inform public health strategies for the development and usage of vaccines and antiviral drugs.

**IMPORTANCE:** We present assays to efficiently sequence genomes of the RSVA and RSVB. This enables researchers and public health agencies to acquire high-quality genomic data using rapid and cost-effective approaches. Genomic data based comparative analysis can be used to conduct surveillance and monitor circulating isolates for efficacy of vaccines and antiviral therapeutics.

## INTRODUCTION

Respiratory Syncytial Virus (RSV) also called human respiratory syncytial virus (hRSV) is a common, contagious airborne viral infection that primarily affects the respiratory tract (1, 2). RSV is a negative-sense enveloped single-stranded non-segmented RNA virus belonging to the *Paramyxoviridae* family, genus *Orthopneumovirus*. This virus has a ~15.2 kb genome that encodes 11 proteins, three surface glycoproteins Fusion protein (F), attachment glycoprotein (G) and the small hydrophobic protein (SH)), RNA dependent RNA polymerase Large protein (L), nucleocapsid (N), phosphoprotein (P), transcriptional regulators (M2-1, M2-2), matrix (M), non-structural proteins (NS1, NS2) (3). The F and G protein promote the production of protective immune response, and antigenic differences in the G, F and SH envelope proteins led to the classification of RSV into two major antigenic group RSV A and B. The F protein is responsible for the fusion of the viral envelope with the host cell membrane for the viral entry into the cell, and is highly conserved and immunogenic which is attractive for vaccine development. In contrast, the G protein, which is responsible for cellular attachment, is prone to frequent mutations (3–5).

RSV diagnosis is usually based on clinical symptoms, but laboratory tests such as rapid antigen tests (6, 7), polymerase chain reaction (PCR) (8) are commonly used diagnostic methods. In recent years, whole genome sequencing (WGS) and its analysis have provided a more in-depth understanding of RSV outbreaks (9–11). By identifying and characterizing viral isolates causing outbreaks, public health authorities and institutions can implement appropriate control measures, such as isolation, contact tracing, and targeted vaccination campaigns (11). WGS-based genomic surveillance involves the systematic monitoring and analysis of viral genomes to understand their genetic diversity, track transmission patterns, and inform public health interventions (12). It has been instrumental in characterizing RSV isolates circulating globally, identifying emerging clades or subclades, and assessing their impact on disease severity and vaccine efficacy. Initially based on G gene sequences, five RSVA and four RSVB clades were identified, named GA1 to GA5 and GB1 to GB5, respectively. Nextclade (https://clades.nextstrain.org/) has now included both G_Clade (13), and recent standardized hRSV Genotyping Consensus Consortium (RGCC) lineages for classifications (14, 15).

WGS based genomic surveillance provides complete genetic profiles on the pathogens and enables researchers to identify specific genetic mutations including single nucleotide polymorphisms (SNPs), insertions, deletions, and rearrangements throughout the viral genomes. Especially since the RNA-dependent replication cycle of RSV is error prone with no proofreading mechanism (16). Analysis of genetic variations and evolutionary patterns informs our understanding of how the virus evolves over time and how new isolates or variants emerge. Furthermore, genomic surveillance helps monitoring genetic changes that may affect the antigenicity of RSV isolates (13). By examining specific genomic regions associated with viral surface proteins (e.g., the G, F and SH), researchers can identify potential variations that might affect the effectiveness of vaccines or antiviral-based therapeutics or prophylactic monoclonal antibodies. This information informs vaccine development strategies and the selection of vaccine isolates (17).

Although there are a number of RSV vaccine candidates in the pipeline (18–20), since late 2023, only two have been approved for use in Canada for adults over 60 years of age including RSVPreF3 (Arexvy) and RSVpreF (Abrysvo™) (20, 21). Both comprise elements of the RSV F stabilized in its pre-fusion conformation (22). Additionally, there are two prophylactic RSV monoclonal antibodies (mAbs) approved for use in Canada including palivizumab (SYNAGIS^®^), which is targeted against the F antigenic site II, and nirsevimab (Beyfourtus™), which is targeted against the F antigenic site Ø. In Canada, the use of palivizumab has been reserved for at risk infants and children under two years of age largely due to cost, limited efficacy, and the requirement of multi-dose regimen. Conversely, nirsevimab, approved for use in Canada in 2023, requires only a single dose and is being recommended for all newborns (22). Nirsevimab makes use of specifically engineered amino acid (aa) changes (M257Y/S259T/T261E) to the fragment crystallizable region of the immunoglobulin G (IgG) antibody that results in an extended half-life in serum capable of providing protection with one injection for the entire RSV season (23, 24). Thus, with wider use of both vaccines and mAbs, it is critical to perform both serological and genomics-based surveillance to identify any shift in antigenicity amongst circulating viruses that could impact RSV vaccines and therapeutics used in Canada (20).

It is worth mentioning that the COVID-19 pandemic has significantly influenced research activities across various fields, including RSV. Although several WGS based methods for RSV have been reported previously, their strategies relied upon amplification of many individual PCR products before pooling them for sequencing (25, 26), hybridization to capture probes, or metagenomic sequencing (27), which are time-consuming and not ideal for high-throughput routine analysis. In this study, we have designed multiplex PCR assays for targeted enrichment to obtain both RSVA and RSVB genomes using Primalscheme (28). For validating our developed multiplex PCR-based WGS methods, and exploring subsequent phylogenetic and mutations analyses, we included a pilot of 89 representative clinical specimens from four different provinces including Ontario (ON), Manitoba (MB), British Columbia (BC) and Saskatchewan (SK) in Canada. Our results demonstrate that the multiplex PCR targeted enrichment method and subsequent genomic analysis have great potential to accelerate large-scale RSV sequencing, diagnosis, and genomic surveillance in Canada. Furthermore, this multiplex PCR assay enables the simultaneous amplification of multiple genomic regions of this virus, allowing for a more comprehensive analysis of its genetic diversity and evolution with enhanced efficiency and sensitivity, which eventually enables researchers to better understand the RSV and inform public health strategies regarding vaccine and antiviral development and usage.

## MATERIALS AND METHODS

### Reference viruses and clinical specimens

Reference RSV isolates used in this study were purchased from Cedarlane and include VR-955 (Strain 9320; originally collected in 1977), VR-1803 (Strain ATCC-2012-11; originally collected in 2012) and VR-1794 (Strain ATCC-2012-10; originally collected in 2012). RSV-positive nasopharyngeal swabs were kindly provided by four provincial laboratories including the British Columbia Centre for Disease Control (BC), Roy Romanow Provincial Laboratory (SK), Cadham Provincial Laboratory (MB), and Kingston Health Sciences Centre (ON). All specimens were collected during the 2022-2023 respiratory virus season with the exception of four specimens from 2016 (SK), one from 2019 (MB) and two from 2021 (BC).

### Development of the Tiled-PCR Amplification Assays with PrimalScheme

#### RSVA

The initial assay was designed against a dataset comprising all publicly available complete RSVA genomes (n=869; accessed November 22^nd^, 2022) downloaded from the Bacterial and Viral Bioinformatics Resource Center (BVBRC) (https://www.bv-brc.org/). The genomes were clustered at 99% sequence identity using cd-hit-est v. 4.8.1 (29) resulting in 71 clusters, and the corresponding genomes representing each cluster were aligned using MAFFT v 7.505 (30). The alignment was re-ordered such that the first sequence corresponded to the largest cluster with subsequent sequences representing smaller clusters. The re-ordered alignment was processed using Primalscheme v. 1.3.2 (28) to develop a multiplex PCR using tiling amplification scheme with default settings, except the amplicon size range was iterated in 100 nucleotides (nts) increments from 400-500 nts through 1100-1200 nts to identify a primer set with the broadest coverage across the genome. The final assay used an amplicon size range of 800-900 nts and the RSVA isolate, SE01A-0167-V02 (Genbank accession: MZ515773.1), was selected as the coordinate system for Primalscheme (28). An additional 2,527 RSVA genomes were acquired from GISAID (accessed November 24^th^, 2022) and combined with those from the BVBRC for a combined dataset comprising 3,396 genomes. After low quality (>10% ambiguous bases) and mislabeled sequences were removed, a total of 3,356 sequences were aligned using MAFFT v 7.505 and the primers output by Primalscheme were mapped against this alignment in Geneious Prime (V2023.2.1, Biomatters Ltd). The primers were manually modified (i.e., shifted up/downstream, application of degenerate nucleotides, or the development of an alternate/supplementary primers) as required to account for any genetic diversity or to correct obvious flaws within the priming region.

#### RSVB

The assay for RSVB was designed similarly as described above for RSVA, but with some changes to simplify the primer design process. The RSVB design was targeted against more contemporary isolates that have been isolated since 2018, which comprised a total of 2,126 partial (≥8000 bases) and complete genomes downloaded from BVBRC (n=437; accessed March 2^nd^, 2023) and GISAID (n=1,692; accessed March 2^nd^, 2023). After low quality (>10% ambiguous bases) and mislabeled sequences were removed, a total of 2,097 sequences were clustered at 99%, 98% and 97% sequence identity using cd-hit-est v. 4.8.1 (29). The set of representative sequences output by cd-hit-est from each sequence identity level were aligned using MAFFT v 7.505 (30), and a consensus sequence was generated from each dataset in Geneious Prime (v2023.2.1, Biomatters Ltd) using the majority nts at each position. The consensus sequences were used as input for Primalscheme v. 1.3.2, and the RSVB isolates, HRSV/B/Bern/2019 (Genbank accession: MT107528.1), was used as the coordinate system to develop a tiled PCR amplification assay using default settings except the amplicon size was set to 500 nts. The primers output by Primalscheme were mapped against an alignment comprising the initial dataset of 2,097 sequences and manually modified as required to account for any genetic diversity or to correct obvious flaws within the priming region.

Prior to ordering, the primers from both assays were re-assessed *in silico* against their respective datasets to identify any primers that required modification (i.e., shifted up/downstream, application of degenerate nucleotides, or the development of an alternate/supplementary primers) to correct for mutations in their extending ends, and/or to account for diversity not captured during the design with PrimalScheme.

### RNA extraction, qRT-PCR and cDNA Synthesis

Viral RNA was extracted from 265 µl of clinical specimens using the Magmax-96 Viral RNA Isolation Kit (Cat No: AMB1836-5, Life Technologies - Invitrogen) as per the manufacturer’s protocol, and the samples were processed on the Thermo Scientific KingFisher Flex Purification System. Viral RNA was eluted in 90 µL of Tris elution buffer and either used immediately or stored at −80°C. The presence of RSV and subtypes was confirmed by quantitative real-time PCR (qRT-PCR) using Invitrogen SuperScript III Platinum One-Step qRT-PCR System (Cat# 11732088) with RSV subtyping primers and probes based on *L* gene (31) and listed here in Table 1. The temperature cycles were one cycle of 50°C for 30 min, one cycle of 95°C for 2 min, 4 cycles of 98°C for 15 s, 63°C for 5 min. SuperScript IV Reverse Transcriptase (Cat#18090200, Invitrogen) was used to synthesize cDNA from 5 µL of RNA in conjunction with 0.5 µL of 60 µM random hexamers [Cat# S1330S, New England Biolabs (NEB)] in a final reaction volume of 10 µL. The reverse transcription reaction was incubated at 42°C for 50 min followed by 70°C for 10 min.

**Table 1:**
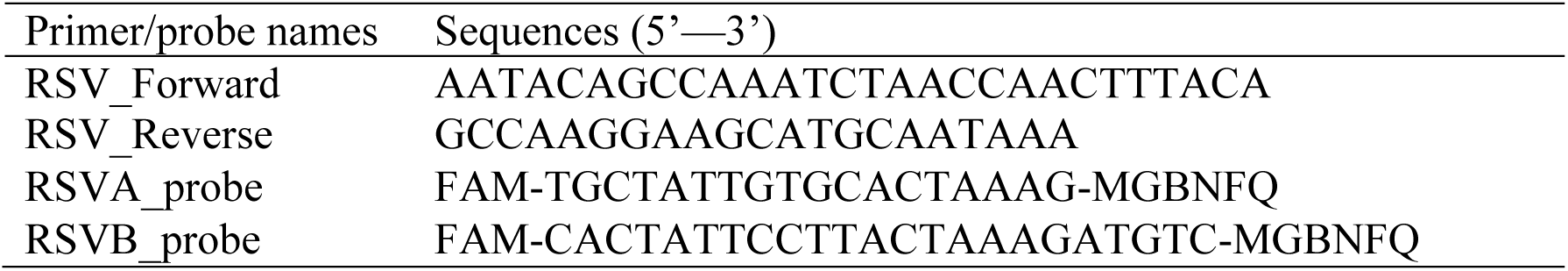
Primer/probe sequences based on L gene used for identifying RSV subtypes.

### Preparation of primer pools and multiplex PCR amplification

For each assay, multiplex primer pools were prepared for each set of primers (i.e., Pool 1 and Pool 2) by combining equal volumes of the appropriate primers (LabReady, 100 uM IDTE, pH 8.0; IDT) then diluting them to 5 µM prior to use. Two separate 25 µL PCR reactions, one for each primer pool, were prepared using the Q5 Hot Start High-Fidelity 2 × Master Mix (Cat# M0494L, NEB) as follows: 12.5 µL of Q5 2 × mastermix, 7.5 µL of nuclease-free water, 2 µL of cDNA and 3 µL of either 5 µM primer Pool 1 or Pool 2. PCR amplification was carried out on MiniAmp™ Plus Thermal Cycler (Applied Biosystems™ by ThermoFisher Scientific) with an initial denaturation stage at 98°C for 30 s, followed by 34 cycles of 98°C for 15 s, 63°C for 5 min, and a final hold at 4°C. The PCR products from each reaction were combined then purified using an equal volume of Ampure XP SPRI Reagent (Cat# A63881, Beckman Coulter) as per the manufacturer’s protocol. The 1 × dsDNA High Sensitivity Kit (Cat# Q33230, Invitrogen) was used to quantify 1 µL of the purified PCR product on the Qubit Flex fluorometer (Invitrogen) as per the manufacturer’s protocol, and the purified PCR products were normalized to 16 ng/µL with 0.01 M Tris in preparation for sequencing.

Each assay was then optimized iteratively against specimens available in-house to identify and correct poorly-amplifying or dropout regions either by modulation of the relative primer concentrations or development of additional replacement primers. The finalized primer sequences and corresponding ratios for both assays are listed in Tables 2 and 3.

**Table 2:**
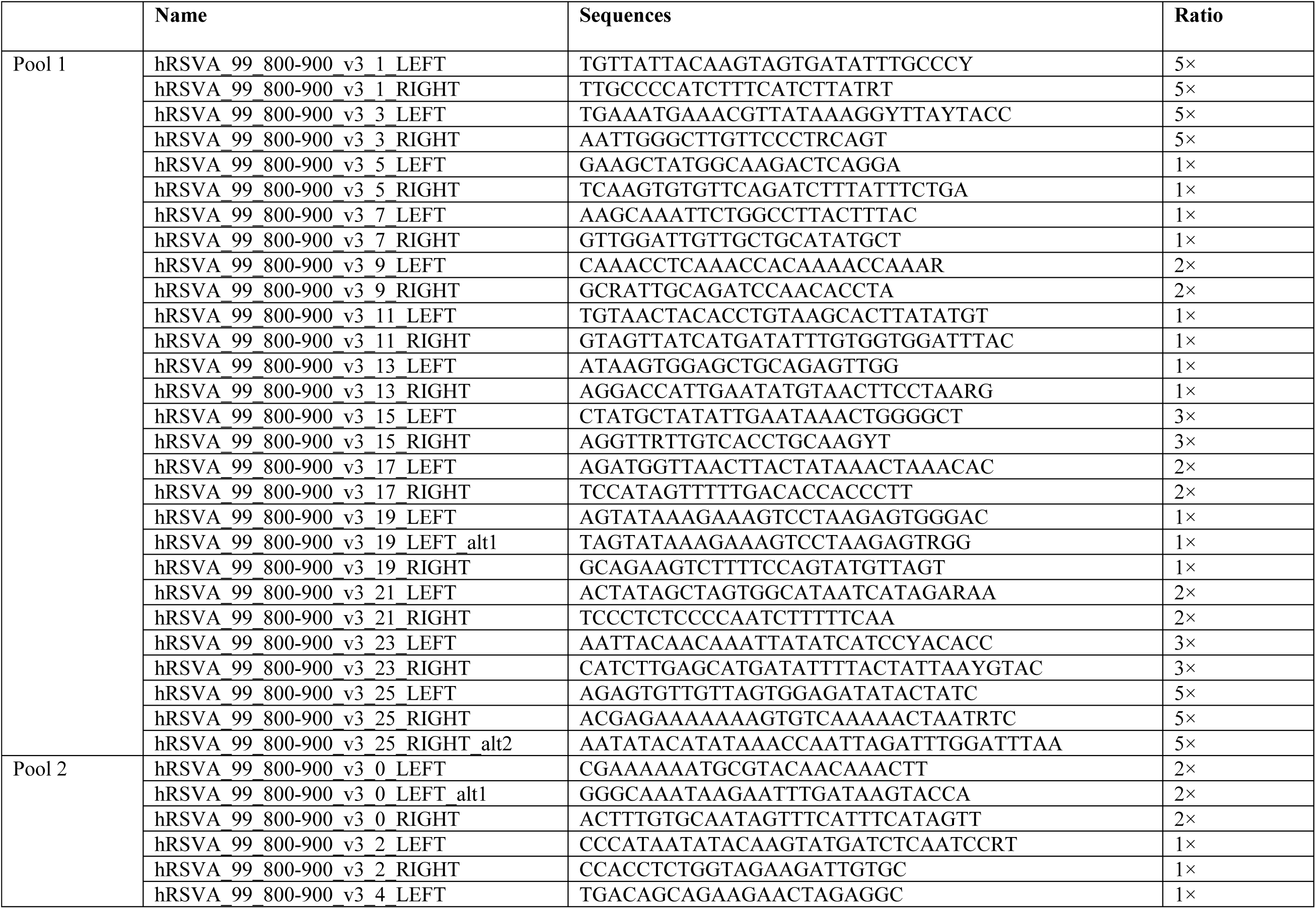
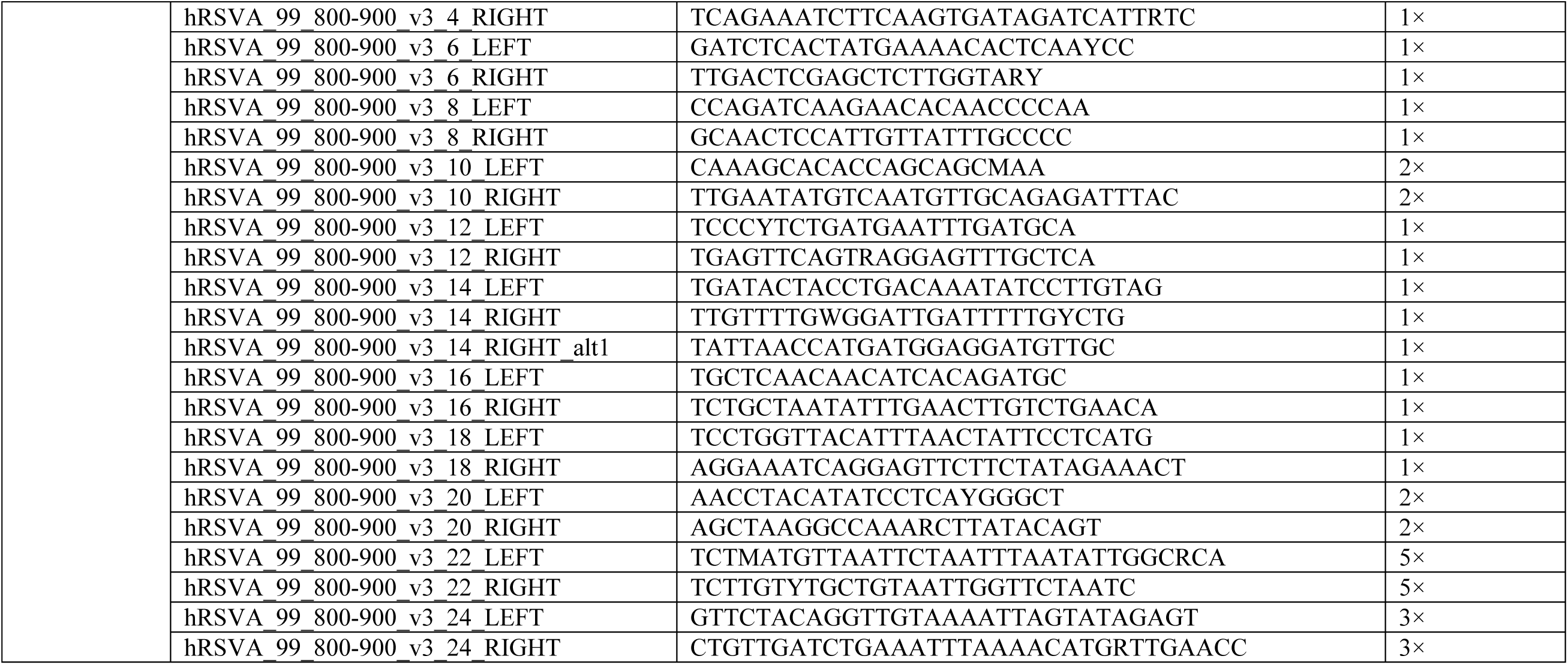
RSVA multiplex PCR primer pools.

**Table 3:**
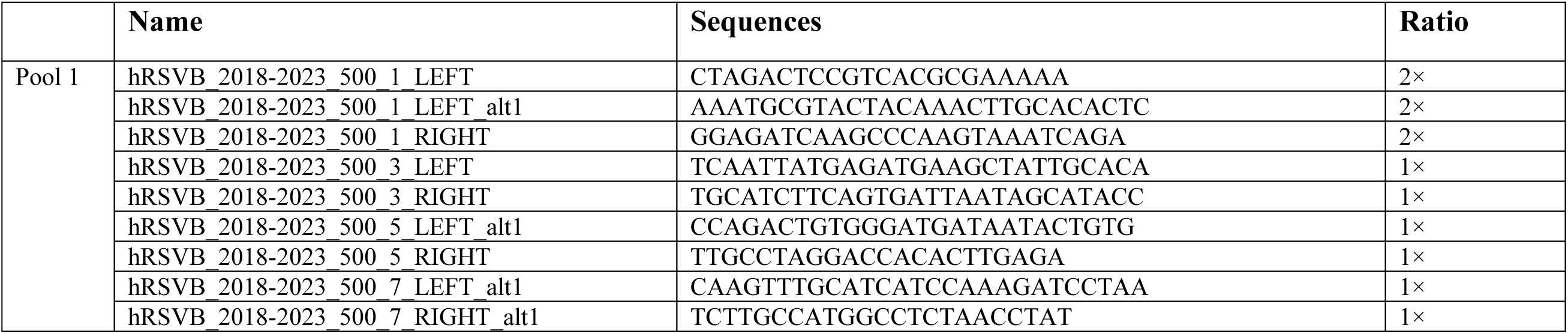
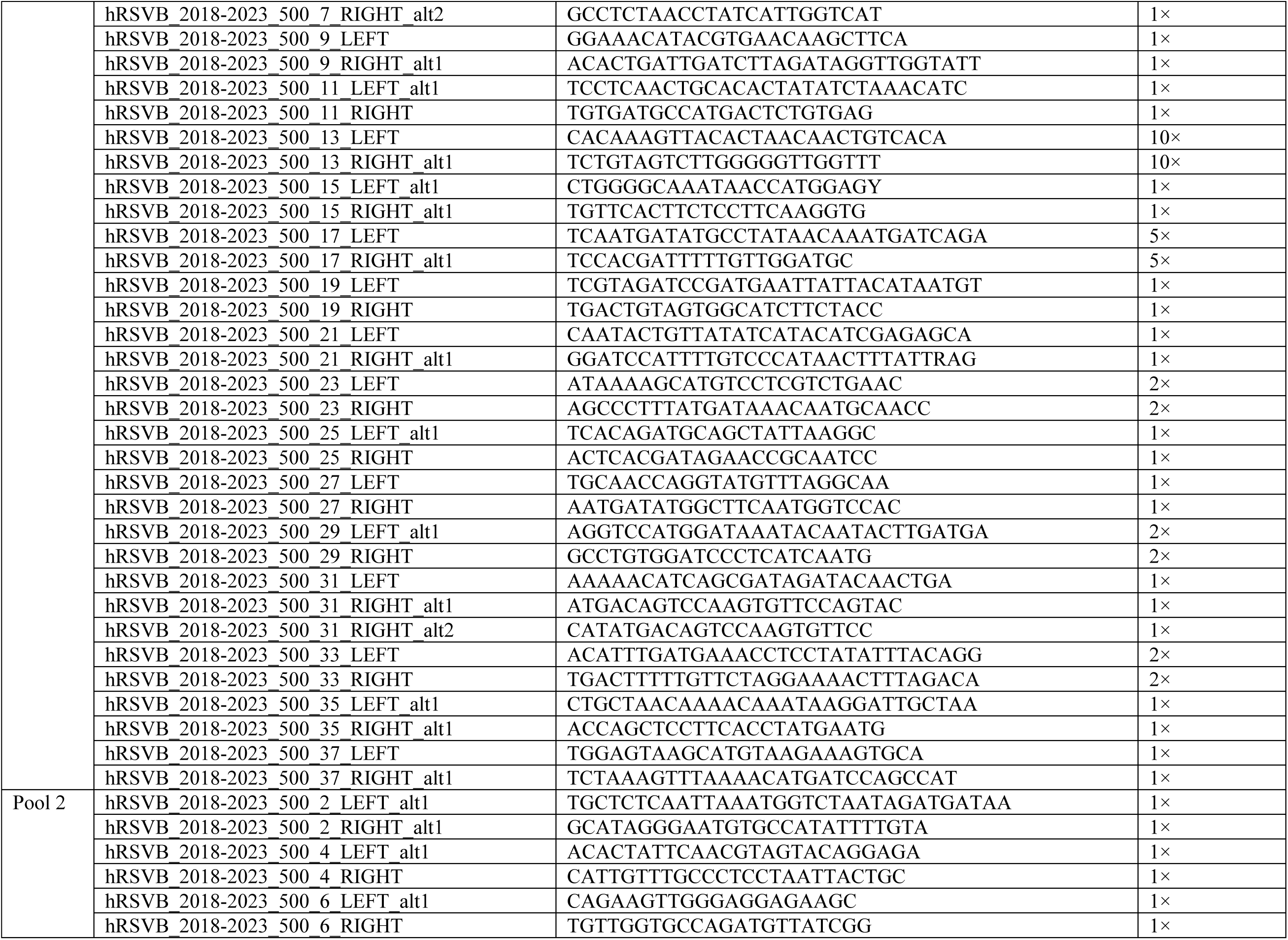
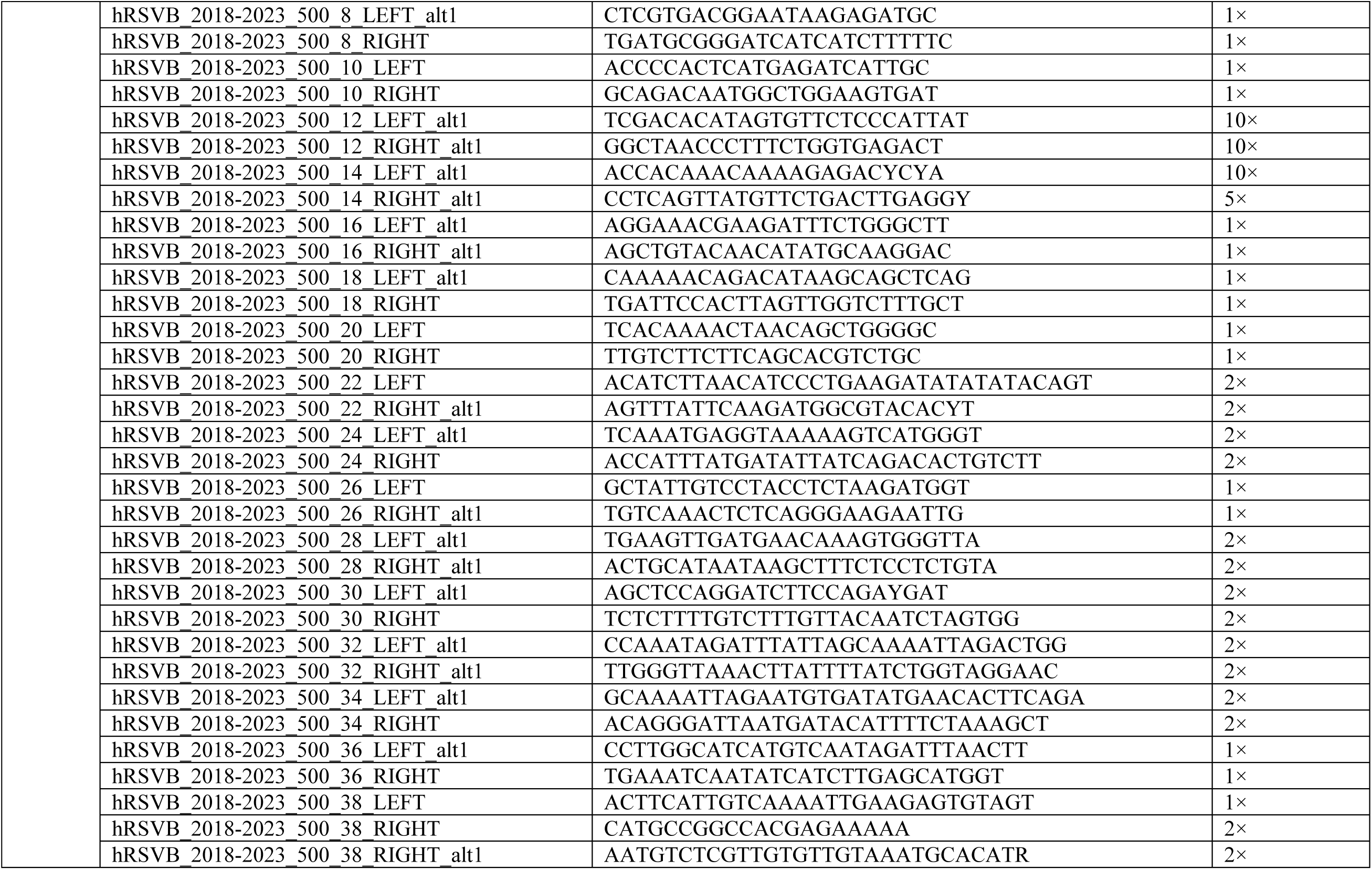
RSVB multiplex PCR primer pools.

### Nanopore library preparation and sequencing workflow

A total of 12.5 µL of normalized PCR product (~200 ng) from each sample was used as input to generate barcoded sequencing libraries using the Native Barcoding Kit (Cat# SQK-NBD114-96, ONT). The protocol was followed exactly with the exception that the NEBNext FFPE DNA Buffer and Repair Mix were not used, and replaced with 1.75 µL of Ultra II End-prep Reaction Buffer. The final library, comprising 12 µL of the pooled libraries, 37.5 µL of Sequencing Buffer and 25.5 µL of Library Beads, was loaded onto a FLO-MIN114 R10.4.1 flow cell with max of 96 specimens and sequenced for 72 h on the MinION Mk1C Sequencing Platform.

### Bioinformatics analysis of sequence data obtained from Nanopore platform

The FAST5 files were basecalled and demultiplexed using Guppy v 6.5.7 with default settings except specification of dna_r10.4.1_e8.2_400bps_5khz_hac as the basecaller model and SQK-NBD114-96 as the barcode kit with the “--require_barcode_both_ends” parameter. The basecalled FASTQ files were processed using the Nextflow-enabled viralassembly pipeline (https://github.com/phac-nml/viralassembly), which is a genericized version of the ncov2019-artic-nf pipeline (https://github.com/connor-lab/ncov2019-artic-nf) that automates the ARTIC Network’s Field Bioinformatics Toolkit (https://github.com/artic-network/fieldbioinformatics). The viralassembly pipeline is capable of processing any tiled-PCR amplification assay as long as the reference sequence (fasta format) and primer coordinate file (bed format) used to create the assay are provided. Briefly, the viralassembly pipeline automates read mapping, primer trimming, variant calling and consensus sequence generation. Default settings were used, except Medaka (https://github.com/nanoporetech/medaka) was selected as the variant caller in conjunction with the r1041_e82_400bps_hac_v4.2.0 model, and the maximum read length to keep was set to 1500 nts for the RSVA assay and 1000 nts for the RSVB assay.

For the purposes of this study, a genome was considered to be complete if it encompassed the first codon of the first NS1 and the last codon of L protein (15). The Open Reading Frames (ORFs) encoding the RSV genes were identified using Geneious Prime (v2023.2.1, Biomatters Ltd) and characterized as a means to detect and correct any sequencing errors, or to validate legitimate biological mutations that would result in the formation of a truncated protein.

### RSVA and RSVB Datasets

In order to contextualize our sequences within the broader population structure of RSV, 124 RSVA and 83 RSVB lineage exemplar reference sequences (https://github.com/rsv-lineages) were downloaded from NCBI. They were combined with the 52 RSVA and 40 RSVB sequences generated here (37 Canadian + 3 ATCC isolates) resulting in datasets comprising 176 RSVA and 123 RSVB sequences, respectively (Supplementary Table 1).

### Characterisation of RSVA and RSVB sequences using Nextclade

The RSVA and RSVB datasets were processed using Nextclade’s webportal on 9 Aug 2024 (https://clades.nextstrain.org/) and analyzed using the RSVA module against the reference isolate hRSV/A/England/397/2017 (EPI_ISL_412866) and the RSVB module against the reference isolate hRSV/B/Australia/VIC-RCH056/2019 (EPI_ISL_1653999), respectively. The RGCC lineage assignments and G_Clade typing nomenclatures (15, 32), as well as quality metrics and genomic features including the genome length, breadth of coverage, and the presence of variants including nts and aa substitutions, insertions and deletions and frameshifts are described in Supplementary Table 1. Amino acid changes identified in the F protein amongst the RSVA and RSVB sequences were tabulated and used to generate heatmaps depicting their carriage across the datasets. The heatmaps were generated using a custom R script implemented in RStudio 2023.12.1 Build 402 running R version 4.3.2 (2023-10-31) with the following packages: ComplexHeatmap v2.10.0, tidyverse v2.0.0, magrittr v2.0.3, grid v4.1.2, and vegan v2.6.4.

### Whole genome phylogenetic analysis

The RSVA and RSVB datasets were aligned separately with MAFFT (v7.520) using default settings (30), then processed with FastTree to infer approximately-maximum-likelihood phylogenetic trees with bootstrap using default settings and the Generalised time reversible (GTR) model (33, 34). The phylogenetic trees were visualized using the Interactive Tree Of Life (iTOL) tool (35) and annotated with the corresponding sequence metadata captured in Supplementary Table 1.

### Glycoprotein G gene phylogenetic analysis

The Glycoprotein (G) ORF for all RSVA and RSVB samples was identified using the “Find ORFs” function in Geneious Prime (v2023.2.1, Biomatters Ltd), and the corresponding coding sequences were extracted and subjected to BLAST analysis to confirm they were in-frame and intact. The G sequences from RSVA and RSVB were each aligned separately using MAFFT (v7.520) with default settings (30), then maximum-likelihood trees were inferred using FastTree with default settings and the GTR model (33, 34). The phylogenetic trees were visualized using the iTOL tool (35) and annotated with the corresponding RGCC lineage and G_Clade assignments from Nextclade, as well as the sample collection year and location (Supplementary Table 1).

### Co-phylogeny analysis

For both RSVA and RSVB, a custom R script was used to construct a co-phylogeny comprising the tree files generated from the whole genome and G gene coding sequences, and annotated with the corresponding RGCC lineage for each sample. The script was implemented as described above using the following packages managed with pacman v0.5.1: ape v5.7.1, biocmanager v1.30.22, tidyverse v2.0.0, ggtree 3.10.1, phangorn v2.11.1, treeio v1.26.0, phytools 2.1.1, viridis, here v1.0.1, and scico v1.5.0.

### Identification of mutations linked to phenotypic or epidemiologic traits using RSVsurver

The RSVsurver, developed by Singapore’s Agency for Science, Technology and Research (A*STAR) Bioinformatics Institute (BII) and enabled by GISAID (https://rsvsurver.bii.a-star.edu.sg/faq.html), was used to screen the RSV sequences against their curated database for the presence of mutations that may be linked to important phenotypic and epidemiological traits. The RSVA and RSVB datasets were uploaded to the RSVsurver and each sequence was compared against the reference isolates hRSV/A/England/397/2017 and hRSV/B/Australia/VIC-RCH056/2019, respectively.

## RESULTS

### Multiplex-PCR based amplification using tiling scheme of RSV and complete genome sequences collection

For RSVA, PrimalScheme generated an assay comprising 24 overlapping amplicons with 0 gaps accounting for 95.5% of the genome. Additional primer sets were manually designed to generate amplicons capturing regions closer to the 5’ and 3’ ends of the genome extending coverage of the assay to >99% of the genome. For RSVB, Primalscheme generated an assay comprising 38 overlapping regions with 0 gaps accounting for 99.6% of the genome. The finalized primer sequences and corresponding ratios for both assays are listed in Tables 2 and 3. Among the in-house 52 RSVA, and 37 RSVB clinical specimens, three ATCC RSVB stains, and Nextclade curated NCBI 124 RSVA and 83 RSVB reference genomes, the antigenic group, subtypes, genome length, identified mutations and GISAID IDs were summarized in the Supplementary Table 1. The near complete genome length ranged from 14,994 nts to 15,225 nts.

### Whole genome phylogenetic analysis of 52 Canadian clinical RSVA isolates

The RSVA dataset (n=176; 52 in-house + 124 reference) was demarcated into nine clades using the G_Clade scheme developed by Goya *et al*., 2020 (Supplementary Table 1 and Fig. 1). The majority of sequences were classified as GA.2.3.5, and largely represent more contemporaneous isolates collected since 2007 including all Canadian isolates sequenced in this study (Fig.1). In contrast, the new scheme proposed by the hRSV RGCC and implemented in Nextclade was more discriminatory and demarcated the same dataset into 25 lineages that more closely reflects the structure of the phylogenetic tree. (Fig. 1; represented by coloured range with the tree nodes). Overall, lineage assignment was well supported by the phylogenetic tree, though not all sequences from lineages A.D and A.D.5 clustered cohesively. However, this was also apparent in the RSVA maximum likelihood tree described in Goya *et al*., 2024 (15) and the publicly available Nextclade RSVA phylogeny.

**FIG 1.**
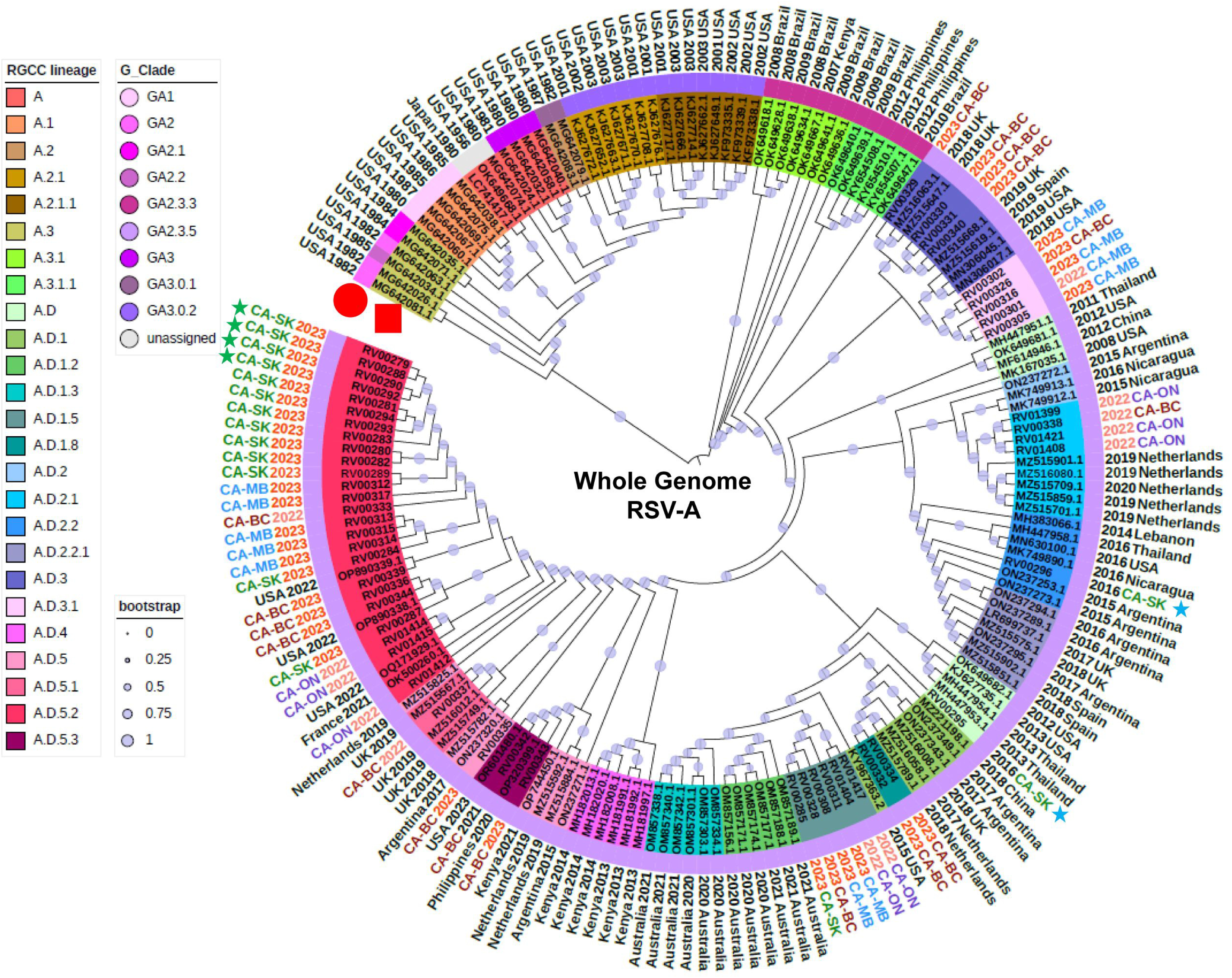
Phylogenetic trees demonstrating 176 RSVA isolates including 52 Canadian isolates and 124 Nextclade references based on whole-genome sequences. hRSV Genotyping Consensus Consortium (RGCC) lineages were shown in different color range and G_Clade was overlaid as a color strip in the phylogenetic tree. The year of collection of 2022 and 2023 were shown in light red and red respectively, and the rest were shown in black. Representative Canadian isolates primarily collected in the year of 2022–2023 and from four provinces, namely British Columbia (BC), Manitoba (MB), Ontario (ON) and Saskatchewan (SK), with the text color of blue, red, purple and green respectively. Bootstrap was annotated in each branch of the tree with light purple dots.

Isolates derived from clinical specimens characterized in this study, collected between 2016 and 2023, were assigned to nine lineages all descending from A.D with the plurality of sequences (25/52) assigned to lineage A.D.5.2 (Fig. 1). The oldest two isolates collected from SK in 2016 were assigned to lineages A.D and AD.2.2, respectively and clustered with reference sequences collected between 2012-2016 (blue stars, Fig.1). Based on available data, lineage A.D.2.2 was prevalent amongst isolates characterized in 2016, but detections have declined since then (15). Interestingly, the remaining Canadian isolates, collected during the 2022-2023 respiratory virus season and one from 2021, were assigned to multiple lineages indicating that a diverse array of viruses can be co-circulating simultaneously. With the exception A.D.5.2 and A.D.1, which represent the most common (n=25) and second most common (n=8) lineages detected amongst the Canadian isolates, a lineage tended to be dominated by sequences from a single province (excluding reference sequences) (Fig.1).

### RSVA glycoprotein G gene based phylogenetic analyses and its co-phylogeny analysis with whole genome-based tree

The structure of the phylogenetic tree derived from the complete coding sequence of the G gene (Fig.2A) was largely congruent with the phylogeny derived from the WGS (Fig.1). Overall, the lineage designations corresponded to well-defined groups in both phylogenies, though use of the WGS showed higher bootstrap values and were better able to resolve the phylogenetic structure within certain sublineages, such as A.D.5.2, and differentiate closely-related sequences that were indistinguishable based on the glycoprotein gene sequence (Fig.1 & Fig.2A). For instance, within the A.D.5.2 lineage, four isolates, RV00279 (CA-SK 2023), RV00288 (CA-SK 2023), RV00290 (CA-SK 2023) and RV00292 (CA-SK 2023) (green stars, Fig. 2A), were indistinguishable using the G gene sequence, but at the WGS level, could be resolved into two pairs of identical sequences: RV00279 (CA-SK 2023) and RV00288 (CA-SK 2023), as well as RV00290 (CA-SK 2023) and RV00292 (CA-SK 2023) (green stars, Fig. 1).

**FIG 2.**
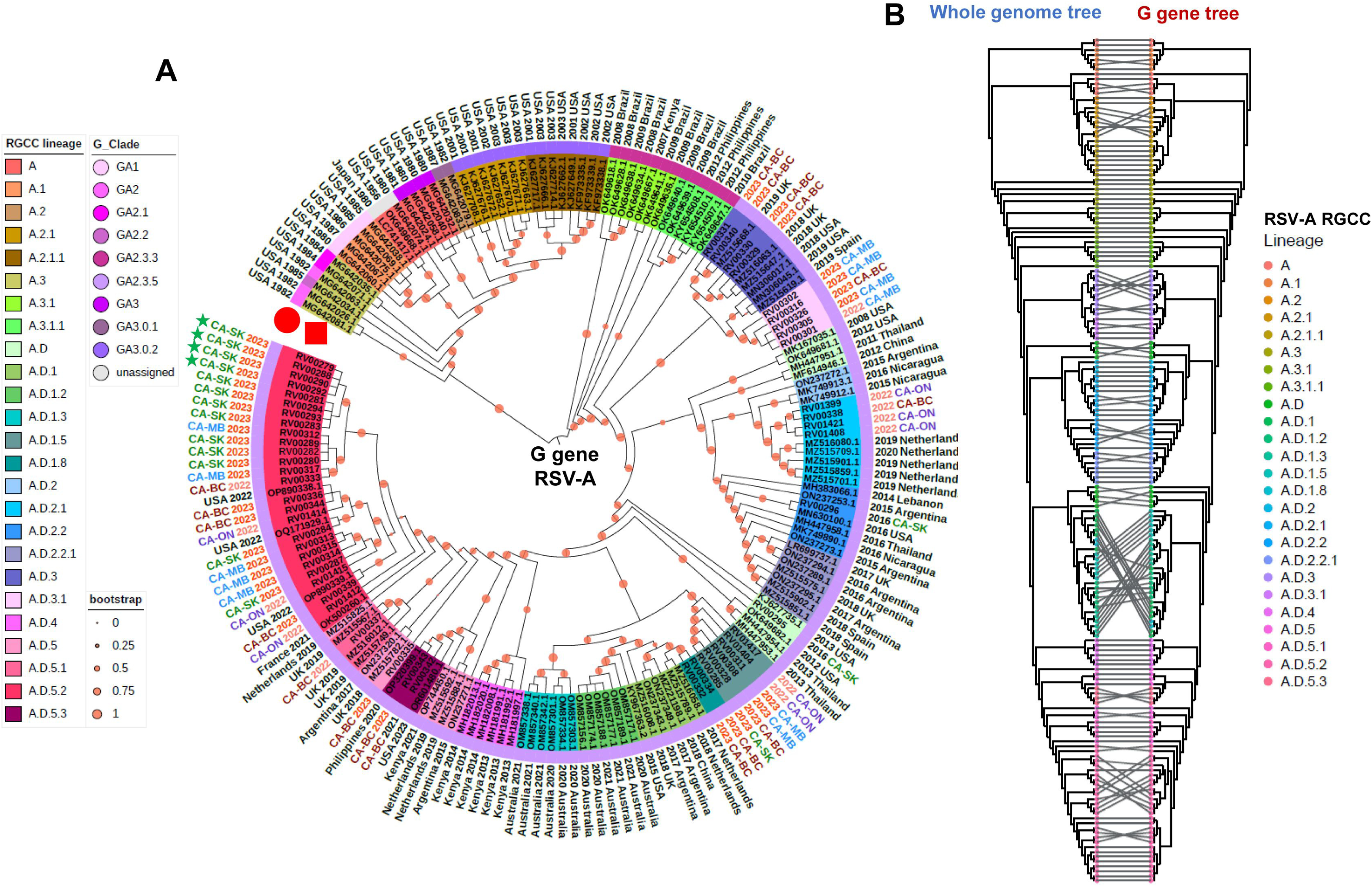
Phylogenetic tree of glycoprotein (G) gene and its comparison with the whole genome-based tree based on 176 RSVA isolates including 52 Canadian isolates and 124 Nextclade references. (**A**) Phylogenetic trees based on G gene sequences. RSV RGCC lineages were shown in different color range and G_Clade was overlaid as a color strip in the phylogenetic tree. The year of collection of 2022 and 2023 were shown in light red and red respectively, and the rest were shown in black. Representative Canadian isolates primarily collected in the year of 2022– 2023 and from four provinces, namely BC, MB, ON and SK, with the text color of blue, red, purple and green respectively. Bootstrap was annotated in each branch of the tree with light red dots. (**B**) Co-phylogenetic tree comparison between whole genome and G gene sequences. RGCC lineages were shown in different colors.

### Whole genome phylogenetic analysis of 37 Canadian clinical RSVB isolates

Using the new RSVB scheme developed by the RGCC, the RSVB dataset (n=123; 37 Canadian + 86 references) were demarcated into 15 lineages that largely reflects the structure of the phylogenetic tree (Fig.3). Comparatively, using the G_Clade scheme developed by Goya *et al*., 2020, the dataset was characterized across seven clades with the majority of sequences classified as GB5.05a, encompassing isolates collected since 2013 including all Canadian isolates sequenced in this study (Fig.3, Supplementary Table 1). One sequence from each of the B (KP856965) and B.D (MH594451) lineages did not cluster cohesively with other sequences from their respective lineages, but this is also reflected in the RSVB maximum likelihood tree described in Goya *et al*., 2024 (15) and the publicly-available Nextclade RSVB phylogeny. Isolates derived from the Canadian clinical specimens were demarcated into 3 lineages with the majority (30/37) assigned to B.D.E.1, which included those from the 2022-2023 respiratory virus season from all provinces, as well as one collected in 2021 from BC. The two isolates collected in 2016 from SK were assigned to B.D.4.1 and clustered with reference sequences collected between 2013-2015, while the remaining three isolates, two collected from MB in 2019 and 2023, and one collected from BC in 2022, were assigned to B.D.4.1.1, which showed a bit further distance as compared to the rest of NCBI reference isolates (Fig.3). The reference viruses VR-955 (collected in 1977), VR-1794 (collected in 2012), and VR-1803 (collected in 2012) were assigned to lineages B, B.D and B.D.4, respectively, and clustered with appropriately contemporaneous reference sequences.

**FIG 3.**
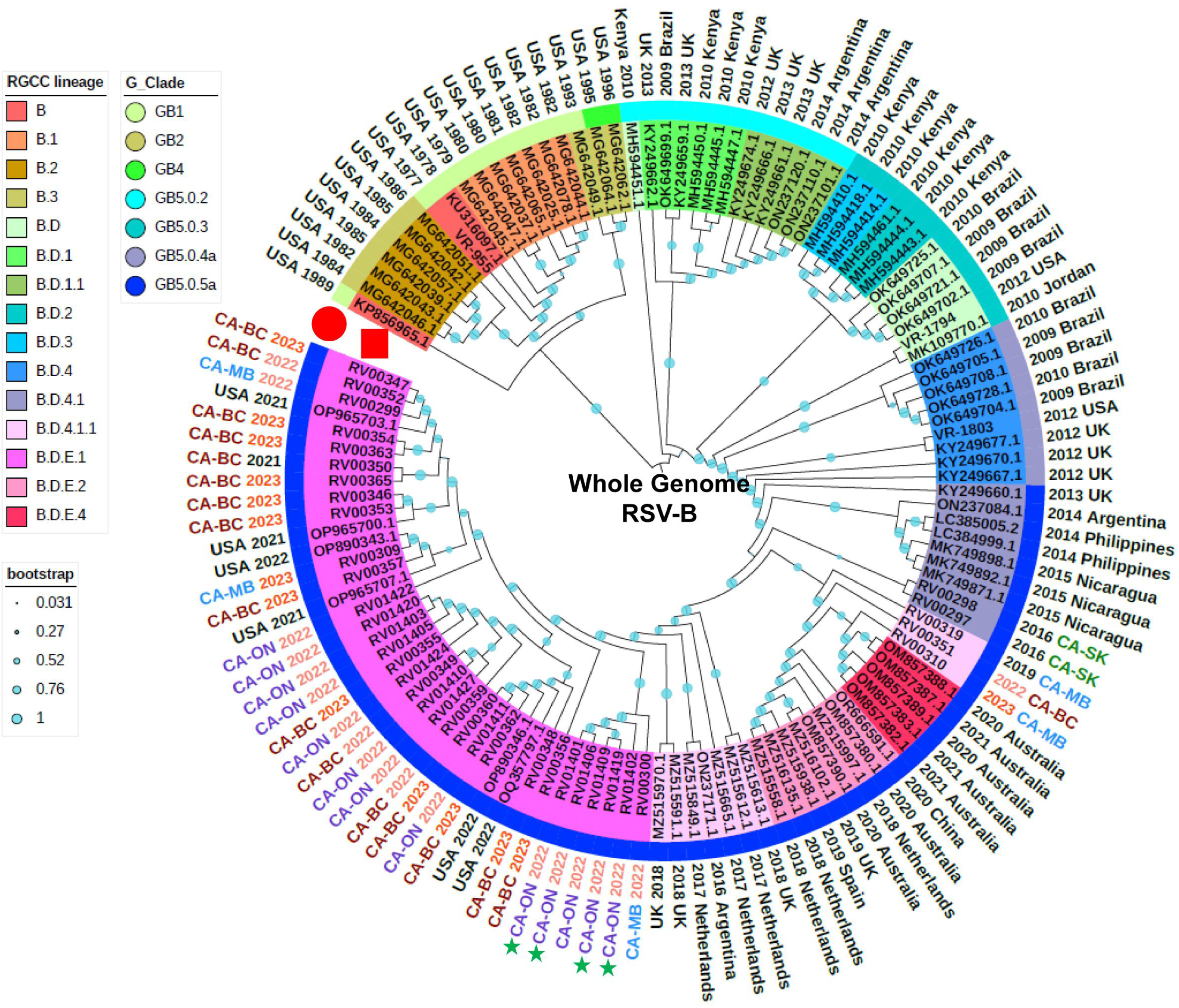
Phylogenetic trees demonstrating 123 RSVB isolates including 37 Canadian and three ATCC isolates, and 86 references based on whole-genome sequences. RGCC lineages were shown in different color range and G_Clade was overlaid as a color strip in the phylogenetic tree. The year of collection of 2022 and 2023 were shown in light red and red respectively, and the rest were shown in black. Representative Canadian isolates primarily collected in the year of 2022–2023 and from four provinces, namely BC, MB, ON and SK, with the text color of blue, red, purple and green respectively. Bootstrap was annotated in each branch of the tree with light cyan dots.

### RSVB glycoprotein G gene based phylogenetic analyses and its co-phylogeny analysis with whole genome-based tree

Similar to RSVA, the structure of the phylogenetic trees derived for RSVB from the complete coding sequences of the G gene (Fig.4A) and WGS (Fig.3) were largely congruent (Fig.4B). Overall, the lineage designations corresponded to well-defined groups in both phylogenies, though use of the complete genome sequences was better able to resolve the phylogenetic structure within certain sublineages such as B.D.E.1, and differentiate closely-related sequences that were indistinguishable based on the glycoprotein gene sequence (Fig.3 & Fig.4A). For instance, in the B.D.E.1 lineage, sequences from four isolates including RV01401 (CA-ON 2022), RV01402 (CA-ON 2022), RV01409 (CA-ON 2022) and RV01419 (CA-ON 2022) (green stars, Fig. 4A) were indistinguishable, but at the whole genome sequence level, could be further resolved into separate clusters (green stars, Fig.3 & 4A).

**FIG 4.**
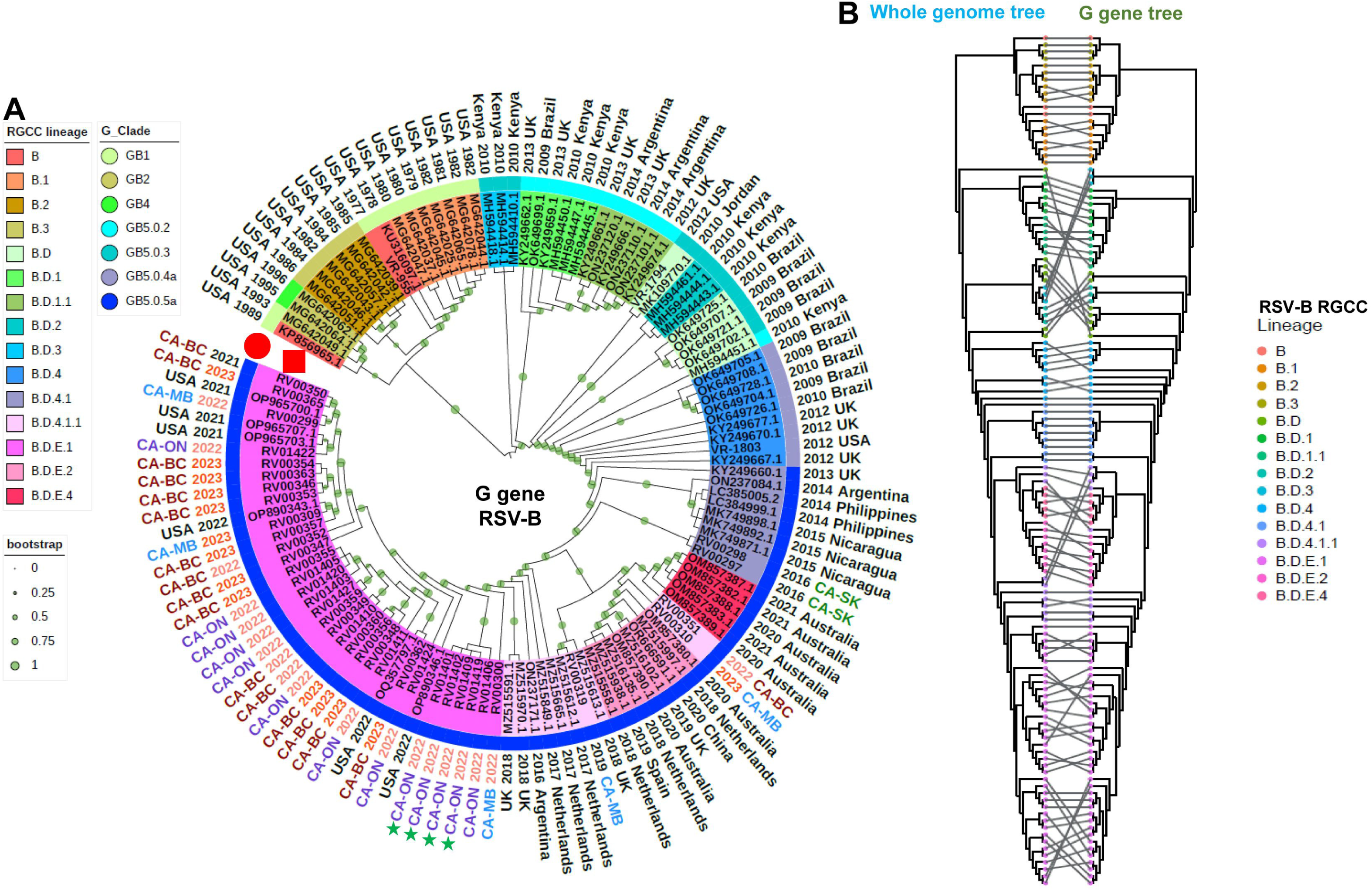
Phylogenetic tree of glycoprotein (G) gene and its comparison with the whole genome-based tree based on 123 RSVB isolates including 37 Canadian isolates and 86 references. (**A**) Phylogenetic trees based on G gene sequences. RGCC lineages were shown in different color range and G_Clade was overlaid as a color strip in the phylogenetic tree. The year of collection of 2022 and 2023 were shown in light red and red respectively, and the rest were shown in black. Representative Canadian isolates primarily collected in the year of 2022–2023 and from four provinces, namely BC, MB, ON and SK, with the text color of blue, red, purple and green respectively. Bootstrap was annotated in each branch of the tree with light green dots. (**B**) Co-phylogenetic tree comparison between whole genome and G gene sequences. RGCC lineages were shown in different colors.

### Characterization of aa substitutions identified amongst the Canadian RSVA and RSVB Sequences

Amongst the 52 Canadian RSVA isolates sequenced in this study, a total of 271 unique aa substitutions were observed across all proteins (NS1=4, NS2=4, N=9, P=11, M=6, SH=4, G=100, F=20, M2-1=14, M2-2=13, and L=86) ranging in frequency from 1.9% to 100% relative to hRSV/A/England/397/2017. Notable mutations flagged by RSVsurver include F:T122A (n=29) and F:T122N (n=2), which negates potential glycosylation of residue 120 (magenta mutations Supplementary Fig. 1), as well as F:K272M (n=1, RV00295, CA-SK 2016, A.D.) and F:S276N (n=5, RV00301-00302, RV00305, RV00316 and RV00326)(Fig. 5A, Supplementary Fig.1). Likewise, for the 37 Canadian RSVB isolates sequenced here, a total of 228 unique aa substitutions were observed across all proteins (NS1=3, NS2=8, N=8, P=6, M=22, SH=4, G=62, F=30, M2-1=4, M2-2=10, and L=71) ranging in frequency from 2.7% to 100% relative to hRSV/B/Australia/VIC-RCH056/2019. RSVsurver flagged the presence of F:R191K in two isolates, RV00297 and RV00298 (CA-SK 2016, B.D.4.1) (orange mutation, Supplementary Fig.1), which was shown to be amongst residues capable of modulating RSV fusion activity *in vitro* (36).

**FIG 5.**
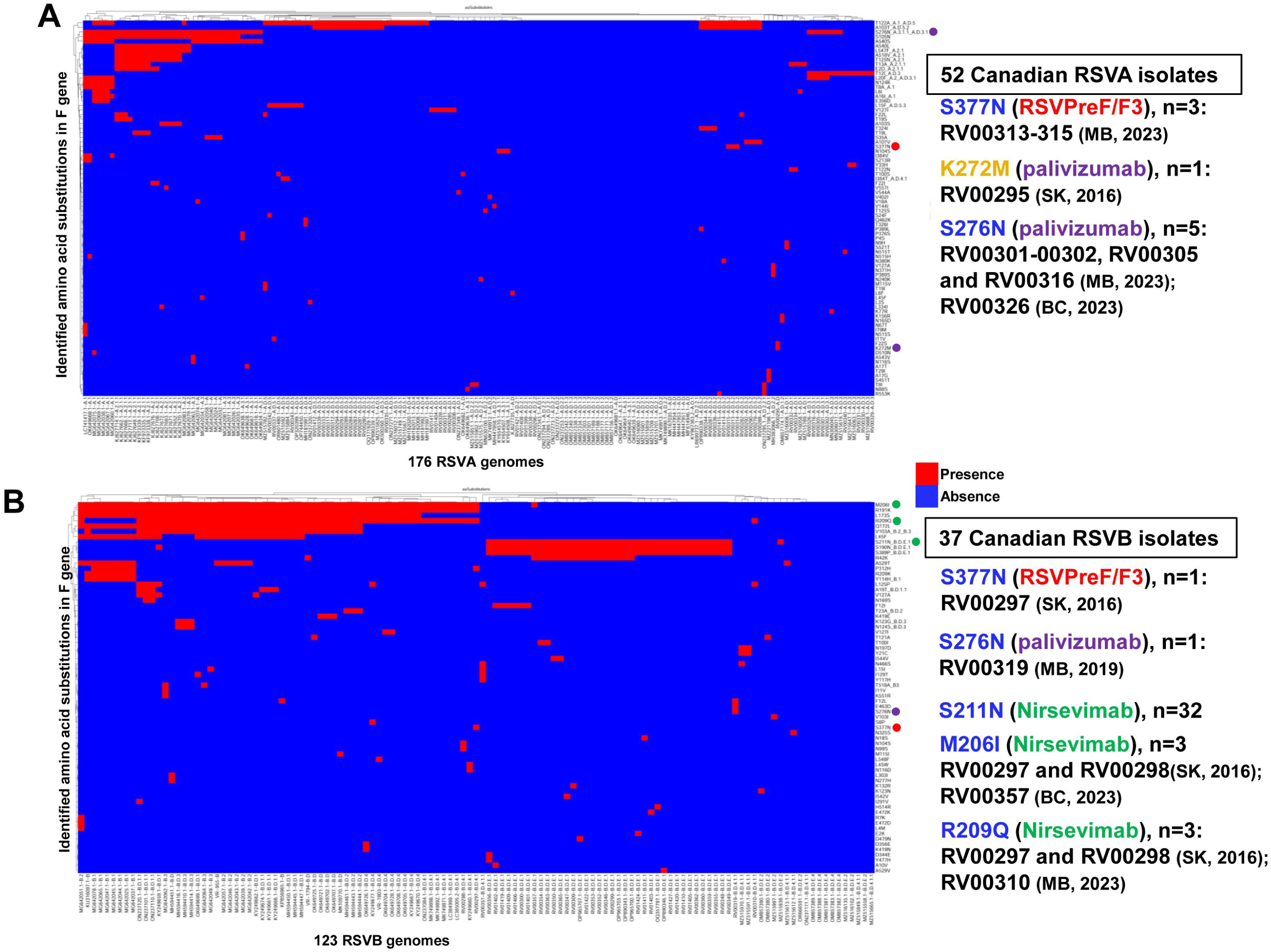
Heatmap distribution of amino acids substitutions in (A) 176 RSVA and (B) 123 RSVB isolates. Red and blue colors represent gene presence and absence, respectively. The x axis contains the list of respective 176 RSVA and 123 RSVB isolates with their lineages indicated and the y axis shows the identified aas substitutions. For 52 Canadian RSVA and 37 RSVB isolates, blue font mutations are assigned interestlevel 1 (moderately significant); orange font mutations are assigned interestlevel 2 (significant), which is known to involved in drug-binding. Vaccines RSVpreF/F3 is in red with red ball pointing to its detected antigenic site mutation. Two mAbs of palivizumab Nirsevimab are shown in purple and green, respectively.

For the Canadian RSVA isolates, no variability was observed within the aa residues corresponding to the nirsevimab targeted antigenic site Ø (62-96, 195-227), while variability was observed at two residues corresponding to the palivizumab targeted antigenic site II (254-277) including F:K272M, as flagged by RSVsurver, as well as F:S276N (n=5); variability was observed at corresponding to the RSVPreF3 targeting antigenic site III (45-54, 301-311, 345-352 and 367-378) including F:S377N (n=3). For RSVB, three residues within antigenic site Ø showed variability including F:M206I (n=3), F:R209Q (n=3), F:S211N (n=32) while F:S276N (n=1) represented the only variability observed within antigenic site II, and F:S377N (n=1) observed within antigenic site III (Fig. 5, Supplementary Figs. 1&2).

## DISCUSSION

The recent licensing of two new RSV vaccines and an additional monoclonal antibody for use in Canada will necessitate implementation of a more robust surveillance system to monitor evolution of this pathogen. To that end, we sought to develop tiled-PCR amplification-based assays for both RSVA and RSVB that can be used to quickly generate near-complete genome sequence data from clinical specimens. This method uses highly-multiplexed PCR amplification to generate overlapping amplicons spanning the targeted genome, and has been used to facilitate sequencing of a variety of different viruses (28, 37, 38). Enrichment by PCR is less expensive and complex, faster, and more portable relative to other methods such as capture probe-based assays, or using techniques like ultracentrifugation and host RNA depletion in combination with brute-force metagenomic sequencing, which is attractive for outbreak investigations, surveillance programs and use in clinical settings. However, the primer design and subsequent multiplex PCR optimization can be challenging, especially for more diverse viral species, and primers may need to be updated over time as the targeted virus evolves (28). With the continued democratization of next-generation sequencing and development of more accessible computational tools for assay design (i.e., Primalscheme), tiled-PCR amplification-based assays are increasingly being developed and deployed to support outbreak investigation and response (Zika), as well as for large-scale genomic epidemiology initiatives (i.e., SARS-CoV-2) (28, 39).

Here, we conducted a small pilot study to test our assays against RSV-positive clinical specimens collected from four different provinces in Canada between 2016-2023. A total of 52 RSVA and 37 RSVB near-complete genomes were recovered and subjected to downstream phylogenetic and comparative genomics analyses. To help contextualize the Canadian sequences within the overall population structure of RSV, we downloaded a collection of lineage exemplar reference sequences and generated phylogenetic trees based on the whole genome and the G gene coding sequences (Supplementary Table 1). Nextclade was used to assign both the new RGCC lineage and G_clade designations for each sequence, and these data were overlaid onto the phylogenetic trees (Figs. 1-4). We observed that the overall structures of the phylogenetic trees derived from the WGS-based and G sequences were largely congruent, though the former was able to distinguish more closely related isolates given the presence of additional sequence data available for interrogation (Figs. 1, 2A, 3 & 4A). This highlights the sensitivity and utility of the whole genome approach to support high resolution outbreak and trace back investigations. Given the contemporaneous nature of the specimens used in this study, all of the RSVA and RSVB sequences contained the A.D and B.D lineage-defining G-gene sequence duplications, respectively. Lineage A.D contains a 72 nts G-gene duplication that emerged in 2011 and by 2017, its descendants had replaced all other lineages (15). Similarly, lineage B.D, first detected in 1999, contains a 60-nt G gene duplication and its descendants replaced all other lineages by 2009 (Supplementary Fig. 3). The evolutionary impact of the duplications in these lineages are not well understood.

From a public health perspective, RSV genomes from different regions and time points provide important information on genetic changes that may affect viral pathogenicity, antigenicity, and vaccine efficacy. This knowledge can ultimately aid in the development of more effective vaccines and antiviral therapies. WGS provides a more complete understanding of the genetic diversity of RSV and permits a comprehensive genomic analysis of specific genomic regions associated with virulence, antigenicity and drug resistance, which is important given the fact that RSV does not employ a proofreading mechanism during replication (16). The assays described here generate near complete genomes that can be readily characterized using both internal and publicly available tools, such as Nextclade and RSVSurver, to facilitate identification of emerging lineages, as well as the presence of biologically important or novel mutations (Supplementary Table 1). RSVsurver was used to visualize the aa changes identified for RSVA and RSVB (colored-balls) among all available antigenic sites within the three-dimensional prefusion and postfusion structure of the RSV F glycoprotein in complex with AM22 (magenta) and Infant Antibody AD-19425 (green) respectively (Supplementary Figs. 1 & 2). The prefusion conformation possesses all six major antigenic sites (Ø, I, II, III, IV, V), of which only I, II, III, and IV are present in the postfusion conformation (40). Analysis using RSVSurver revealed that amongst the Canadian RSVA isolates, one isolate from SK collected in 2016 and three isolates collected from MB in 2023 possess the F:S377N mutation located in antigenic site III targeted by the RSVPreF3 vaccine, and this mutation might be immunodominant sites for both RSVA and RSVB isolates (41). Moreover, one RSVA isolate collected from SK in 2016 contained the F:K272M mutation located in antigenic site II, which has been shown to impact the efficacy of palivizumab (42). Similarly, amongst the Canadian RSVB isolates, three mutations were identified including F:S211N (n=32), F:M206I (n=3) and F:R209Q (n=3) that are located in antigenic site Ø targeted by nirsevimab (Fig. 5, Supplementary Table 1). Thus, these isolates present an opportunity to conduct downstream antigenicity and antiviral testing to study the effect of both the well-characterized and rarer mutations observed across the dataset.

In conclusion, our tiled PCR amplification-based assays represent a convenient and inexpensive, method for the rapid generation of near complete RSV genomes from clinical specimens. These enhanced WGS methods can ultimately contribute to the advancement of RSV research, particularly in the areas of sequencing, diagnosis, and genomic surveillance. We have also demonstrated that the sequence data generated using our assays can be readily analysed using downstream tools including Nextclade for lineage assignment and RSVSurver for screening of important mutations, and can also be used to support epidemiological investigations within a genomic framework. These assays and subsequent genomic analyses offer potential for serving large-scale RSV genomic surveillance with enhanced efficiency and sensitivity, which will allow researchers to better monitor genomic variability in RSV and inform public health strategies for the development and usage of vaccines and antivirals.

## SUPPLEMENTAL MATERIAL

Supplemental material is available online only.

**SUPPLEMENTAL TABLE 1** Profiles and identified mutations among tested 176 RSVA and 123 RSVB genomes

**SUPPLEMENTAL FIG.1** Presence of all amino acid mutations as identified by RSVsurver from the 176 RSVA sequences used in this study contextualized within the 3D structure of the RSV F glycroprotein. (A) prefusion RSVA F glycoprotein (PDB: 6apd, X-ray 4.1 Angstrom) in complex with AM22 (magenta ribbon) & Infant Antibody ADI-19425 (green ribbon). (B) postfusion RSVA F glycoprotein (PDB: 6apb, X-ray 3.0 Angstrom) in complex with Infant Antibody ADI-14359 (green ribbon). The mutations are color-coded by RSVsurver according to their known or predicted biological effect significance. When there are no known effects for the mutation, the mutation will appear in black colored font and assigned interestlevel 0 (least significant). Mutations occurring at a site of interaction will appear in blue colored font and assigned interestlevel 1 (moderately significant). If the mutation occurs at a site known to involved in drug-binding or alters host-cell specificity, it will appear in orange and assigned interestlevel 2 (significant). Mutations will also appear in orange and assigned interest level 2 when its equivalent site is known to result in antigenic shifts or causes mild drug resistance. Mutations that create or remove a potential glycosylation site are colored magenta with assigned interestlevel 2.

**SUPPLEMENTAL FIG. 2** Identified mutations distributed in the different format of 123 RSVB F glycoprotein in complex with antibody. (A) prefusion RSVB F glycoprotein (PDB: 6apd, X-ray 4.1 Angstrom) in complex with AM22 (magenta ribbon) & Infant Antibody ADI-19425 (green ribbon). (B) postfusion RSVB F glycoprotein (PDB: 6apb, X-ray 3.0 Angstrom) in complex with Infant Antibody ADI-14359 (green ribbon). The mutations are color-coded by RSVsurver according to their known or predicted biological effect significance. When there are no known effects for the mutation, the mutation will appear in black colored font and assigned interestlevel 0 (least significant). Mutations occurring at a site of interaction will appear in blue colored font and assigned interestlevel 1 (moderately significant). If the mutation occurs at a site known to involved in drug-binding or alters host-cell specificity, it will appear in orange and assigned interestlevel 2 (significant). Mutations will also appear in orange and assigned interest level 2 when its equivalent site is known to result in antigenic shifts or causes mild drug resistance. Mutations that create or remove a potential glycosylation site are colored magenta with assigned interestlevel 2.

**SUPPLEMENTAL FIG. 3** Duplication to define RSV lineage. (A) Representative sequences demonstration for demonstrating A.D lineage defined by the 72-nt duplication. (B) Representative sequences demonstration for demonstrating A.D lineage defined by the 60-nt duplication.

## DATA AVAILABILITY

All the complete RSV genome sequences were deposited in the GISAID database, with the accession number being listed in Supplementary Table 1.

## ACKNOWLEDGMENTS

This study was funded by Public Health Agency of Canada. We thank Cole Slater from IRVC NMLB for performing wetlab work in this study, Darian Hole from Computational Operational Genomics Section at NMLB, PHAC to develop the viralassembly pipeline (https://github.com/phac-nml/viralassembly), and Ben Hetman for his assistance adapting the R code used to generate the co-phylogeny.

## Notes

The authors declare no conflict of interest.

